# Sex Differences in the Cancer Proteome

**DOI:** 10.1101/2025.03.14.643396

**Authors:** Chenghao Zhu, Nicole Zeltser, Jieun Oh, Constance H. Li, Paul C Boutros

## Abstract

Proteins play a central role in cancer biology: they are the most common drug targets and biomarkers. Sex influences the proteome in many diseases, ranging from neurological to cardiovascular. In cancer, sex is associated with incidence, progression and therapeutic response, as well as characteristics of the tumour genome and transcriptome. The extent to which sex differences impact the cancer proteome remains largely unknown. To fill this gap, we quantified sex differences across 1,590 proteomes from eight cancer types, identifying 901 genes with sex-differential proteins abundance in adenocarcinomas of the lung, and 20 genes across five other tumour types: squamous cell carcinoma of the lung, hepatocellular carcinomas, clear cell cancers of the kidney, adenocarcinomas of the pancreas and glioblastoma. A subset of these protein differences could be rationalized by sex-differential copy number aberrations. Pathway analysis showed that male-biased proteins in lung adenocarcinoma were enriched in MYC and E2F target pathways. These findings highlight the modest impact of sex on the cancer proteome, but the very limited power of existing proteomics cohorts for these analyses.

## Introduction

Proteins play a central role in cancer biology as drug targets and as diagnostic, prognostic and predictive biomarkers – largely quantified *via* immunohistochemistry^1^. Proteomics enables high-throughput profiling protein abundance, structure and modifications, directly quantifying the terminal products of gene expression^2^. The proteome is highly dynamic and is shaped by both genetic and environmental factors. Understanding what determines proteomic characteristics is critical for identifying reliable biomarkers for precision oncology, and in mechanistic studies.

Sex influences gene expression in normal cells. For example, proteomic studies identified thousands of proteins with sex-differential abundance in brain and cerebrospinal fluid, many of which are associated with psychiatric and neurodegenerative conditions^2,3^. Blood proteomic studies revealed substantial sex differences in protein abundance associated with cardiovascular disease and obesity^4,5^. These findings highlight the importance of considering sex as a factor in proteomic studies, and yet the impact of sex on the cancer proteome remains largely unknown.

Indeed, sex has well-documented effects on cancer, with extensive research characterizing sex differences at the genomic level. For example, tumours arising in male patients exhibit higher somatic mutation burdens in cancers such as bladder, melanoma, kidney and liver but lower in glioblastoma^6^. Copy number aberrations (CNAs) differ by sex and are associated with clinical phenotypes^7^. In colon cancer, genes with sex-differential essentiality have been identified^8^, while in pediatric cancers, moderate sex differences in somatic mutations and methylation have been reported (Oh *et al.*, in preparation). These findings suggest sex-specific somatic mutations may at least partially explain sex differences in clinical presentation, including in disease incidence, progression, survival and treatment response^9–15^. While sex differences at the genomic level are well studied, how these differences translate into gene expression, particularly at the protein level, is less understood.

To fill this gap, we analyzed 1,590 high-resolution mass spectrometry-derived proteomes from 934 cancer patients representing eight cancer types. We quantified sex differences in protein abundance in each cancer type using univariable and multivariable statistical modeling, adjusting for key confounders like age and stage. We identified sex differences in protein abundances in six cancer types, with distinct biological pathways influenced by sex. These findings underscore the importance of understanding the influence of sex on cancer molecular profiles, as it can inform patient stratification and guide the development of more effective, tailored treatments.

## Results

### Sex Differences in Protein Abundance

We analyzed 1,590 proteomes across eight cancer types from CPTAC (the Clinical Proteomic Tumour Analysis Consortium). These comprised 915 tumours tissue specimens and 675 normal adjacent tissues (NATs; **Figure 1A**). Cancer types, abbreviations and sample numbers are in **Supplementary Table 1**. The number of samples analyzed ranged from 109 in glioblastoma (GBM) to 314 in liver hepatocellular carcinoma (LIHC)^16–23^. There were 73 ± 21 male and 34 ± 20 female patients per cancer type (median ± MAD; mean absolute deviation). On average, 8,890 ± 637 proteins were detected and quantified per patient. We applied a statistical approach modified from an established two-stage analysis workflow to assess sex differences in protein abundances^6,7,24,25^. Briefly, we first excluded genes with sex differences in normal tissues, followed by univariable analysis to identify putative sex-differential proteins and finally multivariable analysis to control for confounding variables including age and stage (**Figure 1B**; **Supplementary Table 1**).

**Figure 1:**
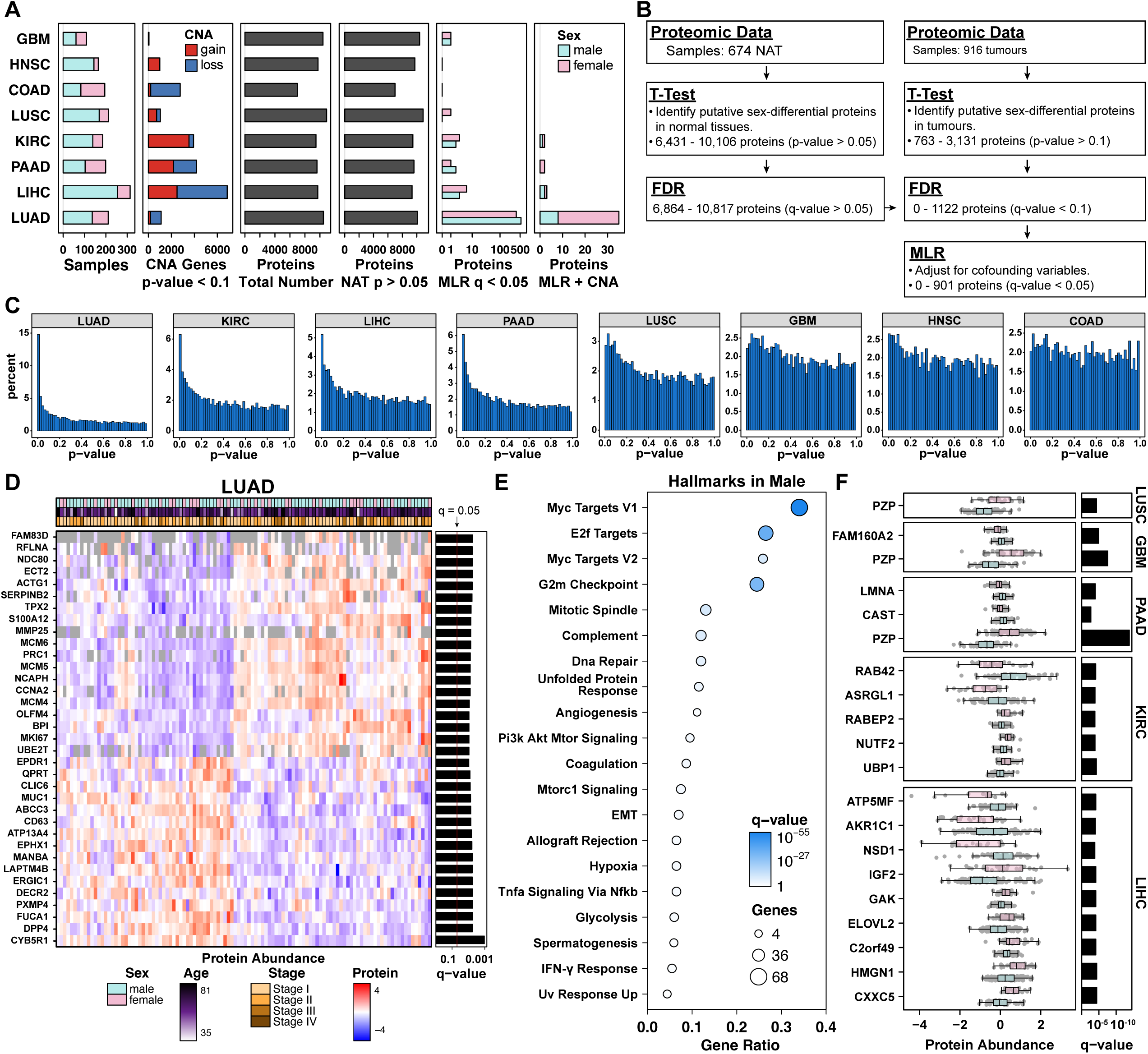
Cancer-type variability in proteomic sex differences **A)** Overview of sex differences in cancer proteome across eight cancer types: lung adenocarcinoma (LUAD), hepatocellular carcinoma (LIHC), clear cell renal cell carcinoma (KIRC), pancreatic and ductal adenocarcinoma (PAAD), lung small cell carcinoma (LUSC), colon adenocarcinoma (COAD), head and neck squamous cell carcinoma (HNSC) and glioblastoma (GBM). From top to bottom, the number of female (pink) and male (blue) patients in each cancer type, total number of proteins quantified and number of genes with significant protein abundant favoring males (blue) and female (pink; q-value < 0.1) in normal adjacent tissues (NATs) and tumours. **B)** Statistical workflow for identifying proteins with sex-differential abundance in tumours. **C)** Distribution of p-values from t-tests comparing protein abundance between sexes in each cancer type. **D)** Heatmap with relative abundance of top 35 genes with significant sex differences in protein abundance (q-value < 0.01, effect size > 0.45 or < −0.45) in LUAD. The top annotation panel indicates patient sex, age and cancer stage. Barplot to the right shows the q-values from MLR. **E)** Top 20 cancer hallmarks enriched among genes with male-biased sex-differential protein abundance in LUAD (q < 0.05). Dot size indicates the number of genes mapped to the each cancer hallmark, while color denotes the q-value. **F)** Boxplots showing genes with significant sex-differential protein abundance (q-value < 0.05, multivariable linear regression (MLR)) in LUSC, GBM, PAAD, KIRC and LIHC. Boxes are colored by sex (blue for male, pink for female). Bar plots to the right show corresponding q-value from MLR.

We first analyzed sex differences in protein abundance in NATs, finding that very few proteins exhibited significant sex-differential abundance (q-value > 0.05; **Figure 1A**; **Supplementary Table 1**), with the exception of lung adenocarcinoma (LUAD) and LIHC, where 369 and 352 proteins showed significant sex difference, respectively; all other cancer types had two or fewer. To assess sex differences in protein abundance of autosomal genes in tumours, we first employed a univariable approach across the eight cancer types after excluding the few proteins that exhibited sex-differences in tissue-matched NAT. We observed pronounced rightward skews in p-value distributions for kidney renal clear cell carcinoma (KIRC), LIHC, LUAD and pancreatic adenocarcinoma (PAAD), indicating strong sex differences in these cancers (**Figure 1C**; **Supplementary** Figure 1). Moderate but clear rightward skews were observed for head and neck squamous cell carcinoma (HNSC) and lung squamous cell carcinoma (LUSC; **Figure 1C**; **Supplementary** Figure 1).

Putative sex-differential proteins (q-value < 0.1, t-test) were further adjusted using multivariable linear regression (MLR) for confounders known to impact gene expression^24,25^, including age, race, smoking, BMI, tumour stage and grade (**Supplementary Table 1**). In LUAD, 901 proteins remained significant (q-value < 0.05, **Supplementary Table 2**), with the top 35 genes (q-value < 0.01 and effect size > 0.45 or < −0.45) shown in **Figure 1D**. Several of these genes have established roles in tumour biology. Among male-biased proteins, *CCNA2* and *MKI67* (q-value = 7.7×10^-3^ and 1.1×10^-3^; effect size = 0.65 and 0.52) are key regulators of cell cycle progression and proliferation, commonly upregulated in LUAD and associated with poor prognosis^26,27^. *PRC1* and *TPX2* (q-value = 7.5×10^-3^ and 3.9×10^-4^; effect size = 0.55 and 0.54) are involved in mitotic spindle organization and chromosomal stability^28–30^, and *SERPINB2* (q-value = 6.0×10^-3^; effect size = 0.47) has been linked to immune modulation and favorable clinical outcomes^31,32^. In contrast, several female-biased proteins such as *EPDR1*, *QPRT* and *ABCC3* (q-value = 9.4×10^-3^, 8.1×10^-4^ and 6.7×10^-4^; effect size = −0.53, −0.48 and −0.86) have been linked to proliferation, metabolic regulation or drug resistance in lung cancer and other cancer types^33–35^. Cancer hallmark enrichment analysis showed genes with significant male-bias are significantly enriched in MYC targets (q-value = 1.0×10^-55^), E2F targets (q-value = 4.2×10^-37^) and G2M checkpoint (q-value = 1.1×10^-32^; **Figure 1E**).

Twenty proteins were identified across LUSC, GBM, PAAD, KIRC and LIHC, with each cohort yielding 1 to 9 significant proteins (q-value < 0.05, **Figure 1F**). No proteins remained significant after MLR adjustment in HNSC and COAD (**Figure 1A**; **Supplementary** Figure 1; **Supplementary Table 2**). Power analyses estimated that at least 187 and 181 samples are needed to detect 20 sex-differential genes in these cohorts (effect size range: > 0.30 or < −0.20 for HNSC and > 0.13 or < −0.12 for COAD; **Supplementary** Figure 2). Thus while our analysis exploits the largest available proteomics cohorts to-date, statistical power to identify sex differences is limited to relatively large effect-sizes.

### Protein Abundance Differences in Genes with Sex-Differential CNAs

Given the variability of sex differences in protein abundance across cancer types and the known influence of CNAs on gene expression, we next focused on genes with sex-differential ploidy status. Using data from the PCAWG (Pan-Cancer Analysis of Whole Genomes) project, we identified 1,939 ± 2,100 genes with putative sex-differential CNAs within each of the eight cancer types (proportion test p-value < 0.1 without adjusting for confounders; **Figure 1A**; **Supplementary Table 3**). Given the high correlation among CNAs across the genome, a strict multiple-testing correction was not applied at this stage to avoid excluding biologically relevant signals. We then integrated CNA findings with proteomic data by comparing genes with sex-differential CNAs within each cancer type to those identified through MLR analysis of sex-differential protein abundance, with multiple-testing adjustment. In PAAD, KIRC, LIHC and LUAD, 42 genes with putative sex-differential CNAs also showed significant (q-value < 0.1) sex-differential protein abundance (**Figure 2A**). In contrast, no sex-differential CNAs showed significant (q-value < 0.1) sex-differential protein abundance in colon adenocarcinoma (COAD), GBM, HNSC or LUSC (**Supplementary Table 2**).

**Figure 2:**
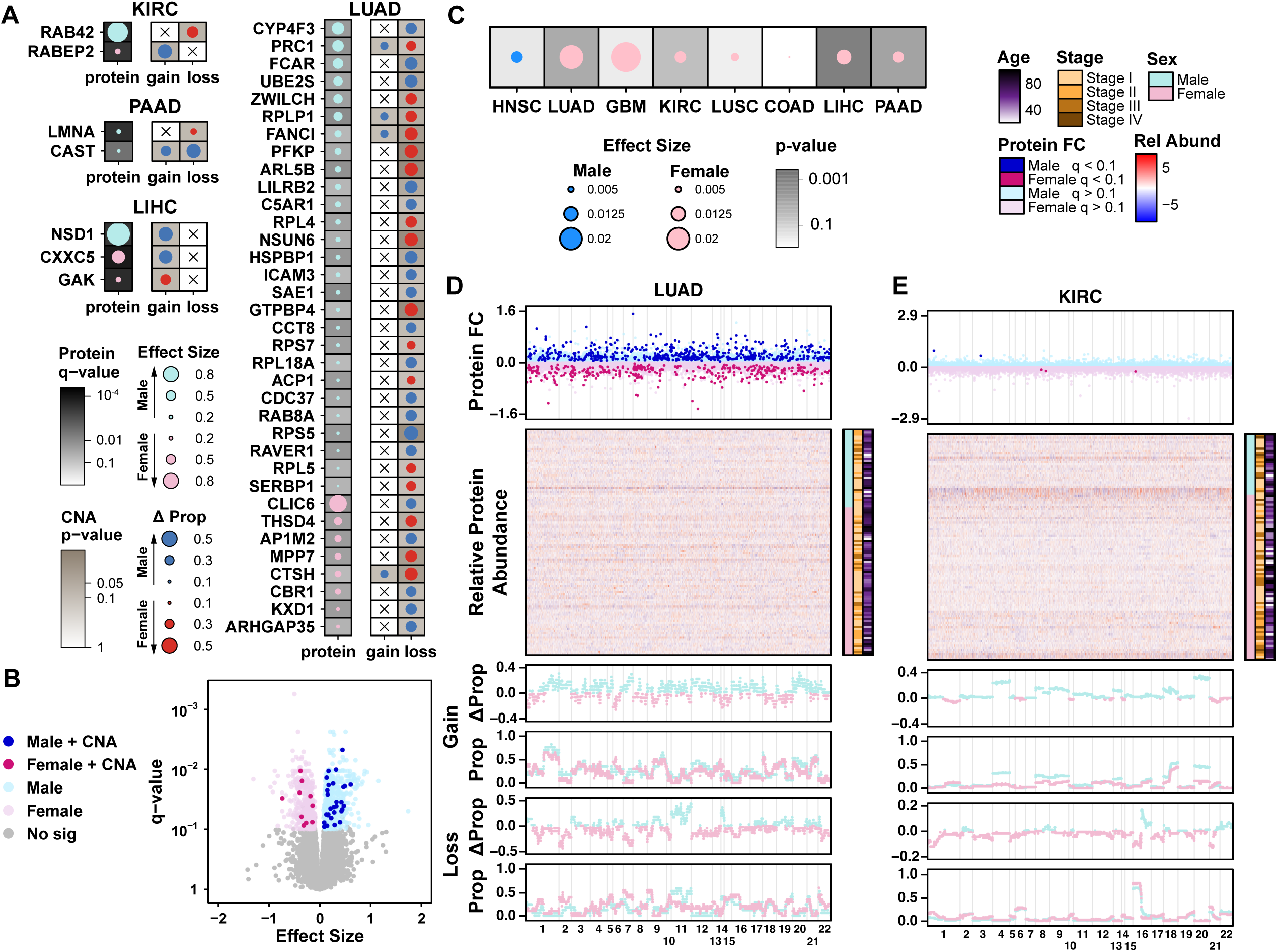
Association between sex differences in protein abundance and ploidy status **A)** Dotmap of genes with significant sex-differential protein abundance and corresponding CNA gain or loss in lung adenocarcinoma (LUAD), clear cell renal cell carcinoma (KIRC), pancreatic ductal adenocarcinoma (PAAD), and hepatocellular carcinoma (LIHC). For each cancer type, the left panel shows multivariable linear regression (MLR) results for protein abundance, and the right panel shows the proportion test results for CNA status. Background shading reflects q-values (protein) or p-values (CNA), dot size represents effect size (protein) or the difference in the proportion of patients with CNA gain/loss, and dot color indicates male (light/dark blue) or female (pink/red) bias. Crosses indicate non-significant CNA differences. **B)** Volcano plots showing effect sizes and q-values for sex difference in protein abundance from tumours derived from LUAD patients. Effect sizes and q-values were calculated using t-test with false discovery rate correction. Light blue and pink indicate a significant bias in protein abundance (q-value < 0.1) favoring males and females, respectively Dark blue and pink represent an additional biased copy number aberrations (CNA) for males and females. **C)** Dot map of Mann-Whitney U test comparing the effect sizes of sex differences in protein abundance between genes with and without sex-differential copy number aberrations (CNAs). Dot color indicates the direction of effect size (blue for male-biased, red for female-biased), dot size represents effect size of Mann-Whitney U test, and background darkness corresponds to statistical significance (darker indicates lower p-values). **D-E)** From top to bottom: fold change (FC) in protein abundance (males - females), heatmap of protein relative abundance, sex differences of copy number gain (males - females), proportion value of copy number gain in males (blue) and females (pink) separately, sex differences of copy number loss (males – females), and proportion value of copy number loss in males (blue) and females (pink). Stacked figures are aligned according to the gene coordinates in the human genome. Stacked panels are aligned according to the gene coordinates at the human genome.

Seven genes with sex-differential protein abundance also had putative sex-differential ploidy change in PAAD, KIRC or LIHC, with no overlap across cancer types (**Figure 2A**; **Supplementary Table 2**). *RAB42* and *RABEP2*, two members of the Ras oncogene superfamily, had opposite sex difference in protein abundance and CNA status in KIRC tumours. *RAB42* had male-biased protein abundance (q-value = 8.2×10^-5^; effect size = 1.02) and higher proportion of female patients with CNA loss (p-value = 0.050; proportion difference (male – female) = −0.12), while *RNBEP2* had female-biased protein abundance (q-value = 1.3×10^-4^; effect size = −0.25) and more male patients with CNA gain (p-value = 0.05, proportion difference (male – female) = 0.15). *CAST*, encoding calpain and prognostic in pancreatic cancer^36^, had elevated protein abundance in males with PAAD (q-value = 2.0 ×10^-3^, effect size = 0.14), and higher proportions of CNA gain (p-value = 0.07, proportion difference (male - female) = 0.07) and loss(p-value = 0.07, proportion difference (male - female) = 0.1) in males. *CXXC5*, a suppressor of the Wnt/β-catenin signaling pathway^37^, showed higher protein abundance in females with LIHC (q-value = 2.7×10^-4^, effect size = −0.73), but higher male patient proportion with CNA gain (p-value = 0.08, proportion difference (male – female) = 0.11).

LUAD tumours had the greatest number of genes with sex-differential protein abundance, however only 35 of the 901 genes showed sex-differential ploidy – thus the majority of sex differences in LUAD were not driven by CNAs (**Figures 2A-B**), none of which overlap with genes significant in PAAD, KIRC or LIHC. Six ribosomal proteins (*e.g., RPS5*, *RPS7, RPL4, RPL5, RPLP1 and PRL18A*) had elevated protein abundance in male LUAD tumours (q-values between 5.7×10^-3^ and 0.033, effect size between 0.11 and 0.42; **Supplementary Table 2**) and varying CNA loss patterns by sex. For example, PRS5 and RPL18A had higher proportion of CNA loss in males (p-values = 0.013 and 0.080, proportion differences (male – female) = 0.34 and 0.43), while RPLP1, RPL4, RPS7 and RPL5 had more CNA losses in females (q-values between 0.043 to 0.083, proportion differences (male – female) between −0.25 to −0.34). *PFKP*, a key regulator of glycolysis and metabolic reprogramming in lung cancer^38^, showed higher protein abundance in males (q-value = 0.023, effect size = 0.31) and higher proportion of CNA loss in females (p-value = 0.011, proportion differences (male – female) = −0.4,), highlighting sex-specific differences in metabolic pathways. *CLIC6*, as mentioned above with increased abundance in females, had more male patients with CNA loss (p-value =0.070, proportion differences (male – female) = 0.31). These findings suggest distinct biological processes, including protein synthesis, metabolism and immune modulation, influenced by sex in LUAD.

### Pan-Cancer Consistency and Molecular Correlates of Sex Differences

To evaluate the consistency of sex-differential protein abundance across cancer types, we examined the correlation of effect sizes between each pair of cancer types (**Supplementary** Figure 3). Overall, effect sizes showed weak to moderate correlations. Although sex-differential protein abundance was identified in PAAD, KIRC, LIHC and LUAD, little correlation was observed with other cancer types. GBM and LUSC, which showed no significant sex-differential protein abundance, exhibited the largest positive correlations, suggesting shared regulatory mechanisms influencing sex-differential protein abundance (Spearman’s Rho = 0.2, p-value = 5.1×10^-88^). These results indicate that sex-differential proteins and the magnitude of differences vary substantially between cancer types.

Given the large number of genes with significant sex-differential protein abundance in LUAD, we next investigated whether these differences were mirrored at the RNA level. We analyzed the transcriptome data from the CPTAC-LUAD project and found a correlation between RNA and protein abundance for the 39 genes with sex-differential protein abundance, as reported above (**Supplementary** Figure 4).

To understand impact of sex-difference in genomic alterations on the cancer proteome, we performed integrated analysis with CNA alterations. Previously, we reported sex differences in CNAs in both coding and noncoding regions of the genome at the pan-cancer level and in specific cancer types, including KIRC and LIHC^6,7^. We applied the Mann-Whitney U test to compare the effect sizes of sex differences in protein abundance between genes with and without sex-differential CNAs. In LUAD, KIRC, PAAD and LIHC, the cancer types where sex-differential proteins were identified, genes with sex-differential CNA status tended to have lower effect sizes in protein abundance, indicating that their protein levels were generally higher in female tumours (p-value from 3.4×10-4 to 0.023; **Figures 2C-E**). In contrast, for the other cancer types, no significant enrichment of sex-differential protein abundance was observed among genes with sex-differential CNAs, suggesting that additional regulatory mechanisms contribute to these differences (**Supplementary** Figure 5).

## Discussion

Our analysis of cancer proteomes revealed variable proteomic sex differences across specific cancer types. Pronounced differences were observed in LUAD, while KIRC, PAAD, LIHC, GBM and LUSC showed moderate variability. These may be driven by interactions between sex hormones (*e.g.*, estrogen and androgen), environmental exposures and the tumour microenvironment, which together shape the sex-dependent evolution of cancers. For example, several P450 enzymes, involved in the metabolism of tobacco-derived carcinogens, are regulated by sex hormones and may further contribute to carcinogenesis^39^. Estrogen and progestin are known to promote angiogenesis in lung cancer^40^, altering the tumour microenvironment. X-inactivation escape in certain X-linked genes may also lead to imbalanced expression of autosomal genes between males and females through regulatory mechanisms^41^. These factors all contribute to sex differences in protein abundance. Interestingly, the male-dominant lung cancer subtype, LUSC, exhibited minimal sex-related differences in the proteome, suggesting distinct tumour origins and progression pathways in LUAD and LUSC. This highlights the differential influence of endogenous and exogenous factors on tumour evolution and gene expression, leading to varying disease manifestations between sexes.

Interpreting sex differences in proteomics is intrinsically complex due to clinico-epidemiological variables partially correlated with sex or influenced by gender-associated behaviors. We applied a statistical approach to identify sex-differential proteins, beginning with exclusion of genes showing sex difference in normal tissues, followed by univariable screening in tumours and multivariable regression to adjust for confounders such as age, race, stage, tumour site, smoking and BMI^6,7^. The impact of unmeasured factors like alcohol consumption, diet, exercise and environmental exposures is a significant limitation. Ideally, sex differences of tumour molecular signatures would be directly quantified using multivariable models with comparisons of their estimates and confidence intervals. The limited sample sizes of cancer proteogenomic cohorts constrains the ability to derive precise estimates, particularly given these numerous confounders.

Indeed while our study extends our understanding of sex differences from genomics to proteomics, it is fundamentally limited by the number of patients and cancer types analyzed. To detect 20 sex-differential proteins with q-value < 0.05 in LUSC, at least 211 patients would be required based on a large effect size threshold of > 1.86 or < −0.22 (**Supplementary** Figure 2). Larger cohorts covering a wider range of cancer types are essential for a more complete picture. Additionally, the datasets we used had imbalanced sample sizes, with fewer female patients in most cancers. Future studies should aim for sex-balanced patient cohorts that reflect the sex-specific prevalence of specific cancer types, ensuring unbiased representation and improving the generalizability of findings. Increasing the sample size is also critical for studying sex differences in low-frequency somatic mutations and their impact on the proteome. To improve statistical power, meta-analysis across independent datasets can be leveraged and statistical methods can help adjust for sample imbalance. Integrating clinical and epidemiological data, including hormone levels, environmental exposures and lifestyle factors, will be critical for understanding the drivers of sex-differential proteomics. While collecting comprehensive metadata is challenging, wearable devices could provide continuous, unbiased tracking of lifestyle variables. Additionally, large-scale biobank data can be leveraged to disentangle the interactions between sex, confounders and tumour proteome alterations. Overall, our findings highlight the importance of investigating sex as a key factor in cancer biology and treatment response, paving the way for personalized therapies based on sex-specific molecular profiles.

## Acknowledgments

This work was supported by the NIH/NCI under awards P30CA016042 and R01CA244729, and by an operating grant from the National Cancer Institute Early Detection Research Network (U01CA214194). CZ and JO were supported by UCLA Jonsson Comprehensive Cancer Center Fellowship Awards. NZ was supported by NHGRI Training Grant in Genomic Analysis and Interpretation T32HG002536 and by NCI training fellowship F31CA281168.

## Author Contributions

**Initiated Study:** PCB, CZ, JO, NZ, CHL

**Data Analysis:** CZ, JO, NZ

**Supervised Research:** PCB, CZ, JO, NZ

**Wrote First Draft of Paper:** CZ, JO, NZ

**Approved Paper:** All authors

## Methods

### Proteomics and Transcriptomics Data

Protein abundance matrices from eight non-reproductive cancer cohorts (HNSC, LUAD, GBM, KIRC, LUSC, COAD, LIHC and PAAD) were obtained from the Clinical Proteomic Tumor Analysis Consortium (CPTAC)^16,17,19–22^. The processed and normalized protein abundance matrices of tumour and normal adjacent tissues (NATs), along with associated clinical information, were downloaded from Proteomic Data Commons (PDC, https://proteomic.datacommons.cancer.gov). Eight samples were excluded from analysis due to failing quality control, either showing high correlation (Pearson’s R > 0.9) or being female samples with high Y-chromosomal protein abundance. Genes allocated on the sex chromosomes are excluded from analysis as they are sex-biased by nature. RNA abundance data (FPKM) from CPTAC’s LUAD cohort were retrieved using the R Package TCGAbiolinks (v3.16).

### Sex Difference of Copy Number Aberrations

Sex difference of copy number aberrations (CNAs) for the eight cancer types were estimated as previously described^6^. PCAWG consensus CNAs (syn8042988) were categorized into gain, neutral, or loss calls per gene. Putative sex-differential genes were identified by comparing the proportions of tumours with gains and losses between sexes using two-tailed proportion tests. No FDR correction was applied at this exploratory stage and genes with a p-value < 0.1 were selected for further analysis.

### Sex Difference of Protein Abundance

A multi-step statistical analysis workflow was used to identify genes with sex-differential protein abundance (**Figure 1B**). First, t-tests were performed on NAT protein abundance to evaluate sex differences, followed by FDR correction (q-value < 0.05). Next, t-tests were applied to tumour samples to assess sex-differential protein abundance, with FDR correction applied (q-value < 0.1). Genes meeting this threshold were further adjusted using multivariable linear regression (MLR) to control for confounders, including age, tumour stage, race and smoking history^6,7^, with FDR correction (**Supplementary Table 1**). MLR results were then annotated with sex-differential CNA status. All statistical tests in this workflow were two-tailed. Cancer hallmark enrichment analysis was performed using the R package hypeR (1.10.0)^42^ and MSigDB gene sets^43,44^ for genes with q-value < 0.1. Mann-Whitney’s test was used to compare the effect sizes of sex differences in protein abundance between genes with and without sex-differential CNAs.

### Sex Differences in RNA Abundance in LUAD

The 39 genes with significant sex-differential protein abundance (q-value < 0.1, MLR analysis) were selected and the corresponding RNA abundances were extracted from the TPM matrix. Multivariable linear regression models were applied to each gene of the tumour samples adjusting confounding variables, including age, tumour stage and smoking history, with FDR correction.

### Power Analysis

Power analysis was conducted per protein for each cancer type, using the effect size and standard error (SE) calculated from t-tests, under the assumption of equal standard errors between males and females and equal numbers of male and female patients. To focus on biologically meaningful differences, we restricted the analysis to the top 20% of genes ranked by absolute effect size in each cancer type. The t-value for each protein was calculated as:

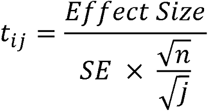

where n represents the total number of samples with cancer type i analyzed in the study, and j is the sample size of interest. For each sample size j, the t values for all proteins were converted into p-values using the degree of freedom j-1, assuming a two-tailed test. These p-values were then adjusted to q-values using the FDR method. The analysis was iteratively repeated across varying sample sizes to determine the number of proteins identified with significant sex differences (q-value < 0.05) at each sample size.

### Statistical Reporting

Results are reported as median values followed by the median absolute deviation (MAD), unless otherwise specified. All boxplots show all data points, the median (center line), upper and lower quartiles (box limits), and whiskers extend to the minimum and maximum values within 1.5 times the interquartile range. All comparisons were performed on biological replicates, defined as independent patients or tumours as appropriate to each analysis.

## Data Availability

Protein abundance matrices were downloaded from Proteomic Data Commons (PDC) with the following accession numbers: PDC000221 (HNSC), PDC000153 (LUAD), PDC000204(GBM), PDC000127(KIRC), PDC000234 (LUSC), PDC000109 (COAD), PDC000198 (LIHC) and PDC000270 (PAAD). RNA-seq data for LUAD are available on Genomic Data Commons (GDC, Project: CPTAC-3, Primary Site: Lung, Primary Diagnosis: Adenocarcinoma, NOS). Data used in this publication were generated by the National Cancer Institute Clinical Proteomic Tumor Analysis Consortium (CPTAC).

## Code Availability

All statistical analysis and data visualization were performed in the R statistical environment (v4.0.2). All visualizations were performed using the BoutrosLab.plotting.general package (v6.0.3) or in Cytoscape.^45,46^ Code is available at https://github.com/uclahs-cds/project-CancerBiology-SexDifferencesv3PedProtFunc.

## References

1. Passaro, A. et al. Cancer biomarkers: Emerging trends and clinical implications for personalized treatment. Cell 187, 1617–1635 (2024).

2. Seo, D. et al. Sex and aging signatures of proteomics in human cerebrospinal fluid identify distinct clusters linked to neurodegeneration. medRxiv 2024.06.18.24309102 (2024) doi:10.1101/2024.06.18.24309102.

3. Wingo, A. P. et al. Sex differences in brain protein expression and disease. Nat Med 29, 2224–2232 (2023).

4. Halland, H., Vitorino, R., Gerdts, E., Meyer, K. & Kararigas, G. Sex-biased proteomic regulation in obesity. European Heart Journal 45, ehae666.2894 (2024).

5. Liu, Y. et al. Sex differences in proteomics of cardiovascular disease: results from the Yale-CMD registry. European Heart Journal 45, ehae666.3091 (2024).

6. Li, C. H. et al. Sex differences in oncogenic mutational processes. Nat Commun 11, 4330 (2020).

7. Li, C. H., Haider, S., Shiah, Y.-J., Thai, K. & Boutros, P. C. Sex Differences in Cancer Driver Genes and Biomarkers. Cancer Res 78, 5527–5537 (2018).

8. Zeltser, N., Zhu, C., Oh, J., Li, C. H. & Boutros, P. C. Sex Differences in Cancer Functional Genomics: Gene Dependency and Drug Sensitivity. 2025.02.05.636540 Preprint at 10.1101/2025.02.05.636540 (2025).

9. Siegel, R. L., Miller, K. D., Fuchs, H. E. & Jemal, A. Cancer Statistics, 2021. CA Cancer J Clin 71, 7–33 (2021).

10. Rahbari, R., Zhang, L. & Kebebew, E. Thyroid cancer gender disparity. Future Oncol 6, 1771–1779 (2010).

11. Konstantinidis, I. T. Trends in Presentation and Survival for Gallbladder Cancer During a Period of More Than 4 Decades: A Single-Institution Experience. Arch Surg 144, 441 (2009).

12. Wainer, Z., et al. Sex-Dependent Staging in Non–Small-Cell Lung Cancer; Analysis of the Effect of Sex Differences in the Eighth Edition of the Tumor, Node, Metastases Staging System. Clin Lung Cancer 19, e933–e944 (2018).

13. Kammula, A. V., Schäffer, A. A., Rajagopal, P. S., Kurzrock, R. & Ruppin, E. Outcome differences by sex in oncology clinical trials. Nat Commun 15, 2608 (2024).

14. Conforti, F. et al. Cancer immunotherapy efficacy and patients’ sex: a systematic review and meta-analysis. Lancet Oncol 19, 737–746 (2018).

15. Milano, G. et al. Influence of sex and age on fluorouracil clearance. J Clin Oncol 10, 1171– 1175 (1992).

16. Huang, C. et al. Proteogenomic insights into the biology and treatment of HPV-negative head and neck squamous cell carcinoma. Cancer Cell (2021) doi:10.1016/j.ccell.2020.12.007.

17. Gillette, M. A. et al. Proteogenomic Characterization Reveals Therapeutic Vulnerabilities in Lung Adenocarcinoma. Cell 182, 200–225.e35 (2020).

18. Satpathy, S. et al. A proteogenomic portrait of lung squamous cell carcinoma. Cell 184, 4348–4371.e40 (2021).

19. Wang, L.-B. et al. Proteogenomic and metabolomic characterization of human glioblastoma. Cancer Cell (2021) doi:10.1016/j.ccell.2021.01.006.

20. Clark, D. J. et al. Integrated Proteogenomic Characterization of Clear Cell Renal Cell Carcinoma. Cell 179, 964–983.e31 (2019).

21. Vasaikar, S. et al. Proteogenomic Analysis of Human Colon Cancer Reveals New Therapeutic Opportunities. Cell 177, 1035–1049.e19 (2019).

22. Gao, Q. et al. Integrated Proteogenomic Characterization of HBV-Related Hepatocellular Carcinoma. Cell 179, 561–577.e22 (2019).

23. Cao, L. et al. Proteogenomic characterization of pancreatic ductal adenocarcinoma. Cell 184, 5031–5052.e26 (2021).

24. Li, C. H., Haider, S. & Boutros, P. C. Ancestry Influences on the Molecular Presentation of Tumours. 2020.08.02.233528 Preprint at 10.1101/2020.08.02.233528 (2020).

25. Li, C. H., Haider, S. & Boutros, P. C. Age influences on the molecular presentation of tumours. Nat Commun 13, 208 (2022).

26. Martin, B. et al. Ki-67 expression and patients survival in lung cancer: systematic review of the literature with meta-analysis. Br J Cancer 91, 2018–2025 (2004).

27. Zhang, L., Wang, S. & Wang, L. Comprehensive analysis pinpoints CCNA2 as a prognostic and immunological biomarker in non-small cell lung cancer. BMC Pulm Med 25, 14 (2025).

28. van Gijn, S. E. et al. TPX2/Aurora kinase A signaling as a potential therapeutic target in genomically unstable cancer cells. Oncogene 38, 852–867 (2019).

29. Hu, J., He, Q., Tian, T., Chang, N. & Qian, L. Transmission of Exosomal TPX2 Promotes Metastasis and Resistance of NSCLC Cells to Docetaxel. Onco Targets Ther 16, 197–210 (2023).

30. Zhan, P. et al. PRC1 contributes to tumorigenesis of lung adenocarcinoma in association with the Wnt/β-catenin signaling pathway. Mol Cancer 16, 108 (2017).

31. Schroder, W. A. et al. SerpinB2 inhibits migration and promotes a resolution phase signature in large peritoneal macrophages. Sci Rep 9, 12421 (2019).

32. Ramnefjell, M., Aamelfot, C., Helgeland, L. & Akslen, L. A. Low expression of SerpinB2 is associated with reduced survival in lung adenocarcinomas. Oncotarget 8, 90706–90718 (2017).

33. Gimeno-Valiente, F. et al. EPDR1 up-regulation in human colorectal cancer is related to staging and favours cell proliferation and invasiveness. Sci Rep 10, 3723 (2020).

34. Zhao, Y. et al. ABCC3 as a marker for multidrug resistance in non-small cell lung cancer. Sci Rep 3, 3120 (2013).

35. Yan, Y. et al. A comprehensive analysis of the role of QPRT in breast cancer. Sci Rep 13, 15414 (2023).

36. Uhlen, M. et al. Towards a knowledge-based Human Protein Atlas. Nat Biotechnol 28, 1248–1250 (2010).

37. An, Z., Wang, J., Li, C. & Tang, C. Signal integrator function of CXXC5 in Cancer. Cell Commun Signal 23, 25 (2025).

38. Shen, J. et al. PFKP is highly expressed in lung cancer and regulates glucose metabolism. Cell Oncol (Dordr) 43, 617–629 (2020).

39. Penaloza, C. G. et al. Sex-dependent regulation of cytochrome P450 family members Cyp1a1, Cyp2e1, and Cyp7b1 by methylation of DNA. FASEB J 28, 966–977 (2014).

40. Marquez-Garban, D. C. et al. Progesterone and estrogen receptor expression and activity in human non-small cell lung cancer. Steroids 76, 910–920 (2011).

41. San Roman, A. K., et al. The human Y and inactive X chromosomes similarly modulate autosomal gene expression. Cell Genom 4, 100462 (2024).

42. Federico, A. & Monti, S. hypeR: an R package for geneset enrichment workflows. Bioinformatics 36, 1307–1308 (2020).

43. Subramanian, A. et al. Gene set enrichment analysis: a knowledge-based approach for interpreting genome-wide expression profiles. Proc Natl Acad Sci U S A 102, 15545–15550 (2005).

44. Liberzon, A. et al. Molecular signatures database (MSigDB) 3.0. Bioinformatics 27, 1739–1740 (2011).

45. P’ng, C. et al. BPG: Seamless, automated and interactive visualization of scientific data. BMC Bioinformatics 20, 42 (2019).

46. Shannon, P. et al. Cytoscape: A Software Environment for Integrated Models of Biomolecular Interaction Networks. Genome Res. 13, 2498–2504 (2003).

