## Supplementary Information for "Sex Differences in the Cancer Proteome"

### Supplementary Figures

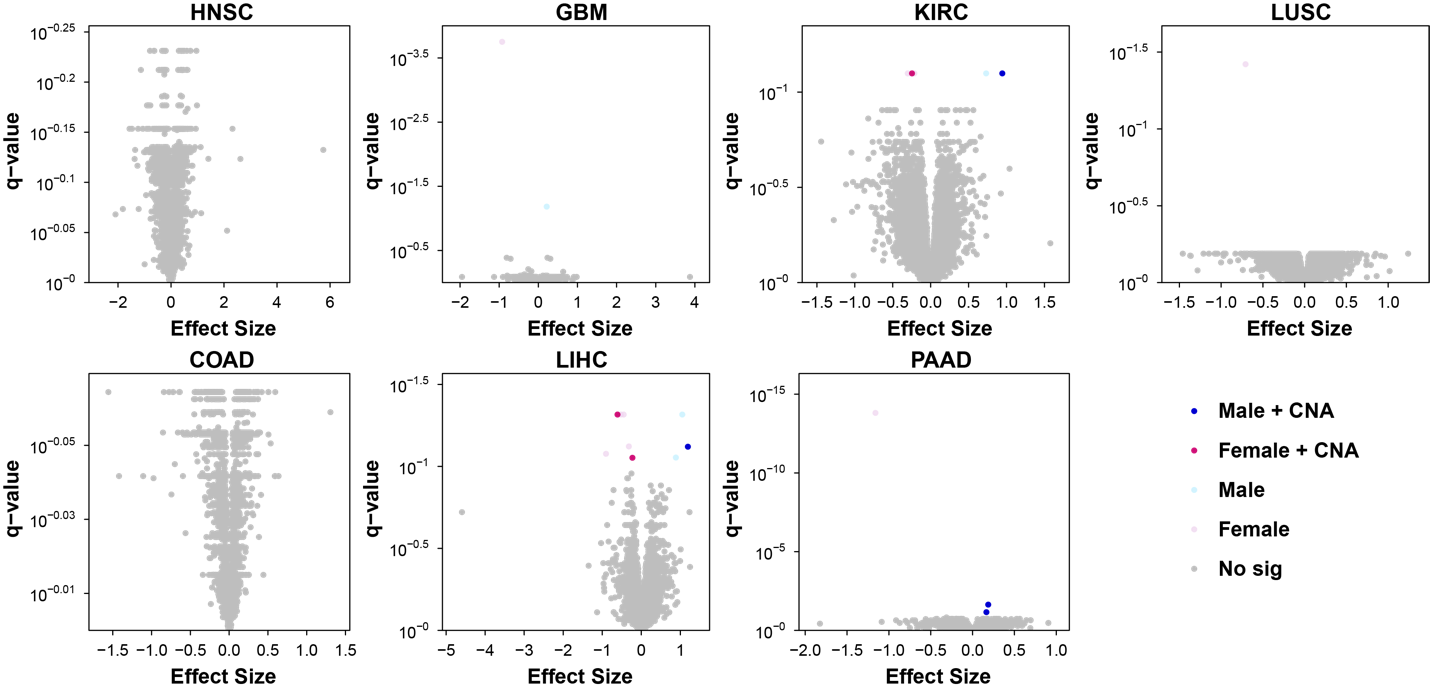

#### Supplementary Figure 1: Volcano plots of Sex Differences in Cancer Proteome

Volcano plots showing effect sizes and p-values for sex difference in protein abundance across eight cancer types: head and neck squamous cell carcinoma (HNSC), glioblastoma (GBM), clear cell renal carcinoma (KIRC), lung squamous cell carcinoma (LUSC), colon cancer (COAD), hepatocellular carcinoma (LIHC) and pancreatic ductal adenocarcinoma (PAAD). Effect sizes and q-values were calculated using t-test with false discovery rate correction. Light blue and pink indicate a significant bias in protein abundance (q-value < 0.1, multivariable linear regression) favoring males and females, respectively, while dark blue and pink represent an additional biased copy number aberrations (CNA) for males and females.

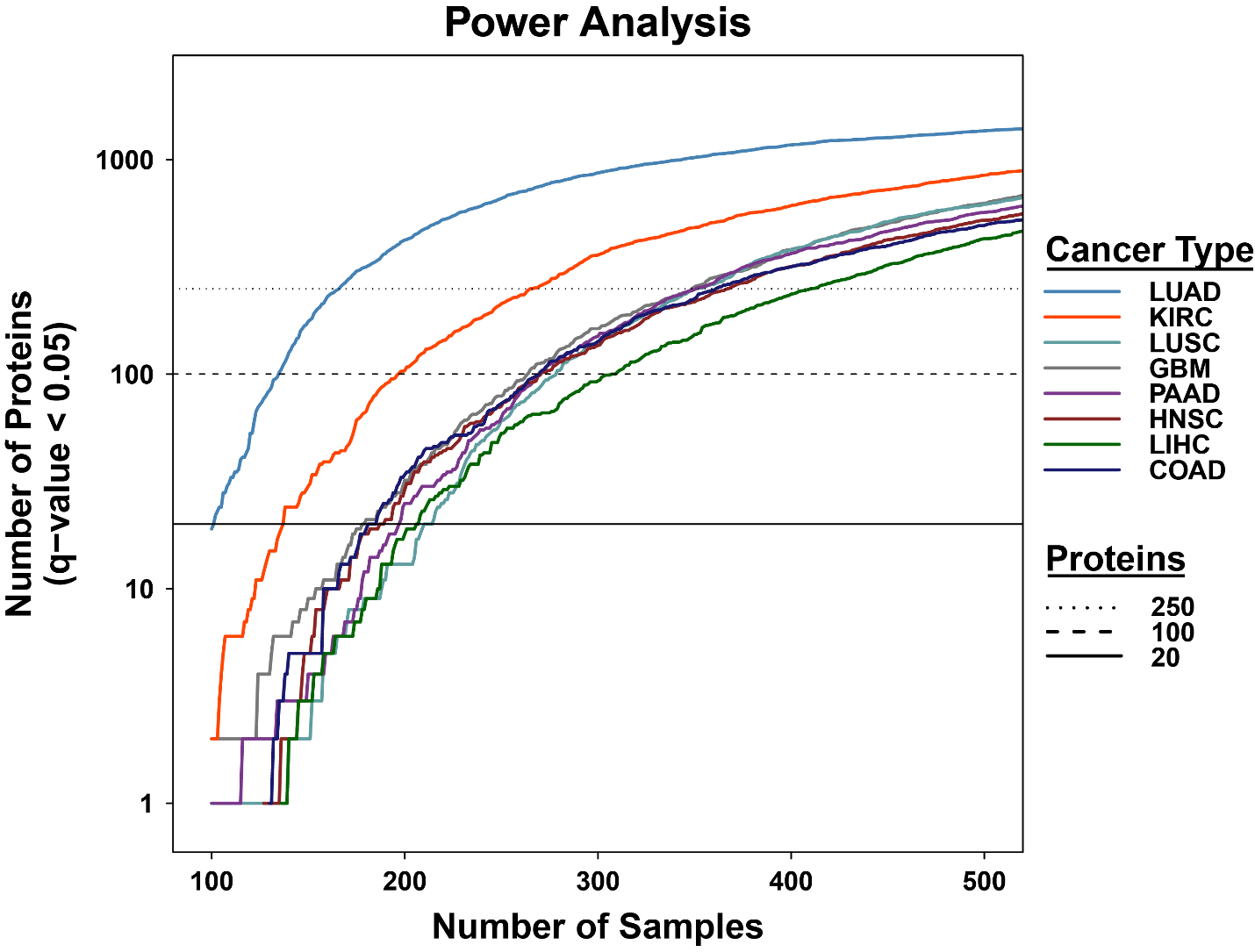
**Supplementary Figure 2: Power Analysis of Proteomic Sex Differences Across Eight Cancer Types.**

The number of proteins detected with significant sex differences (q-value < 0.05) is plotted against the sample size required for each cancer type. Horizontal lines represent thresholds for detecting 20, 100 and 250 sex-differential proteins. Cancer types include lung adenocarcinoma (LUAD), clear cell renal cell carcinoma (KIRC), lung squamous cell carcinoma (LUSC), glioblastoma (GBM), pancreatic ductal adenocarcinoma (PAAD), head and neck squamous cell carcinoma (HNSC), hepatocellular carcinoma (LIHC) and colon adenocarcinoma (COAD).

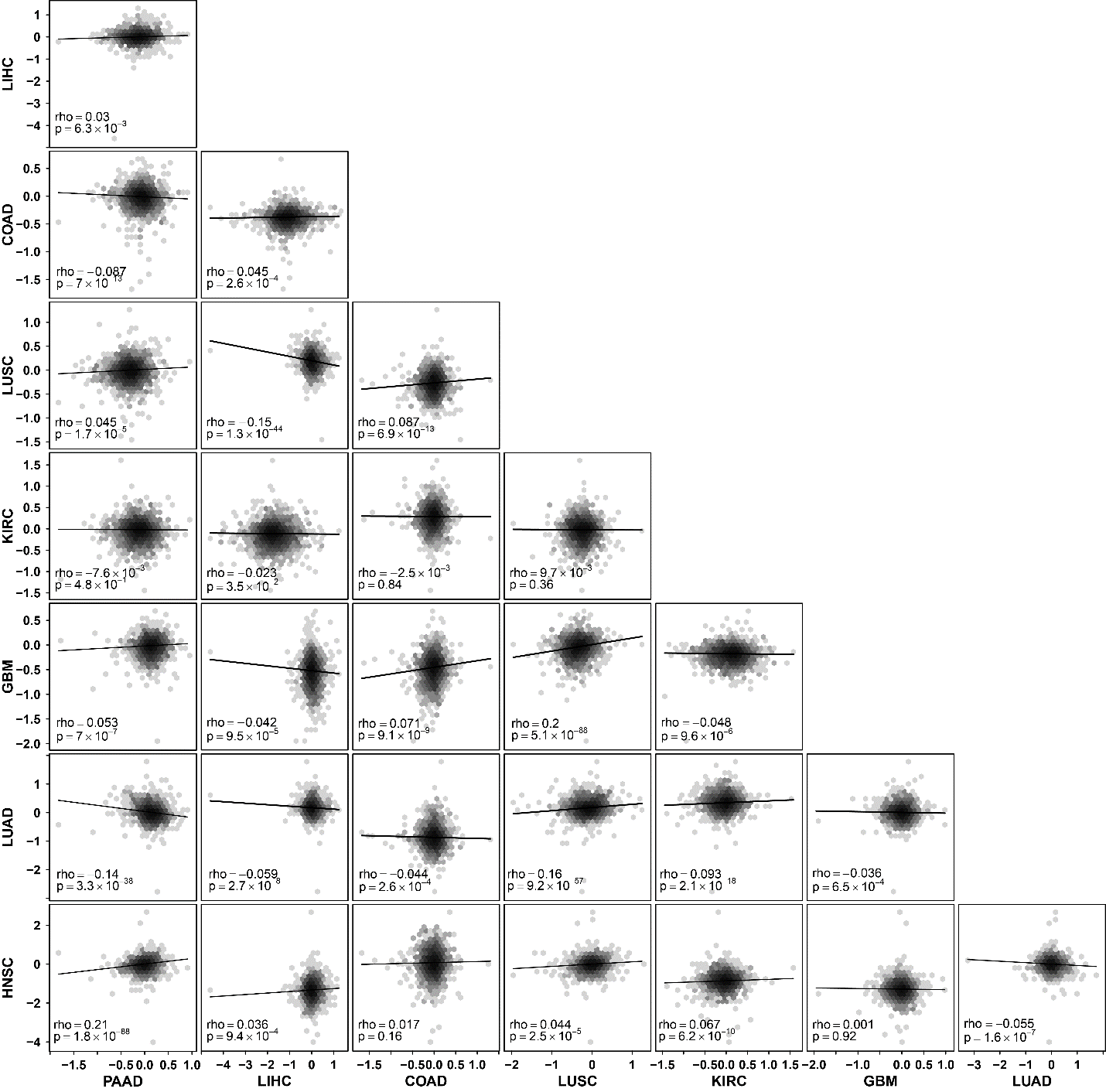

#### Supplementary Figure 3: Correlation of Effect Sizes for Sex-Differential Protein Abundance Across Cancer Types

Hexagonal binned plots show the correlation of effect sizes for sex-differential protein abundance between each pair of cancer types. Each point represents a gene, with the x- and y-axes indicating effect sizes in the respective cancer types. Spearman’s correlation coefficients (rho) and p-values are provided for each comparison.

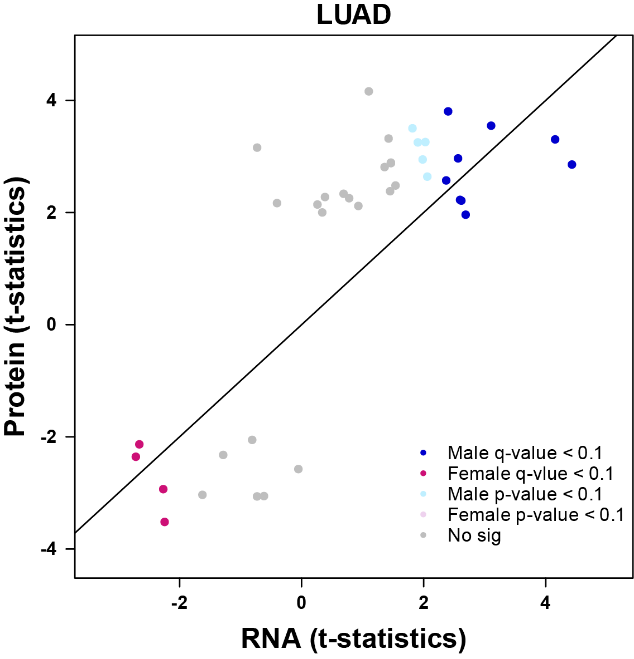

#### Supplementary Figure 4: Correlation of Sex Differences Between Protein and RNA

Correlation between sex differences in protein and RNA abundance in lung adenocarcinoma (LUAD). A multivariable linear regression model was used to calculate the t-statistics for sex difference in protein abundance, while t-test was used for RNA abundance. Dark blue and pink represent genes with significant (q-value < 0.1) sex differences in RNA abundance favoring males and females, respectively, while light blue and pink indicate genes with p-value < 0.1. All genes plotted had significant sex differences in protein abundance (q-value < 0.1). The line in the scatter plot represents equality (y = x), not the best-fitted line.

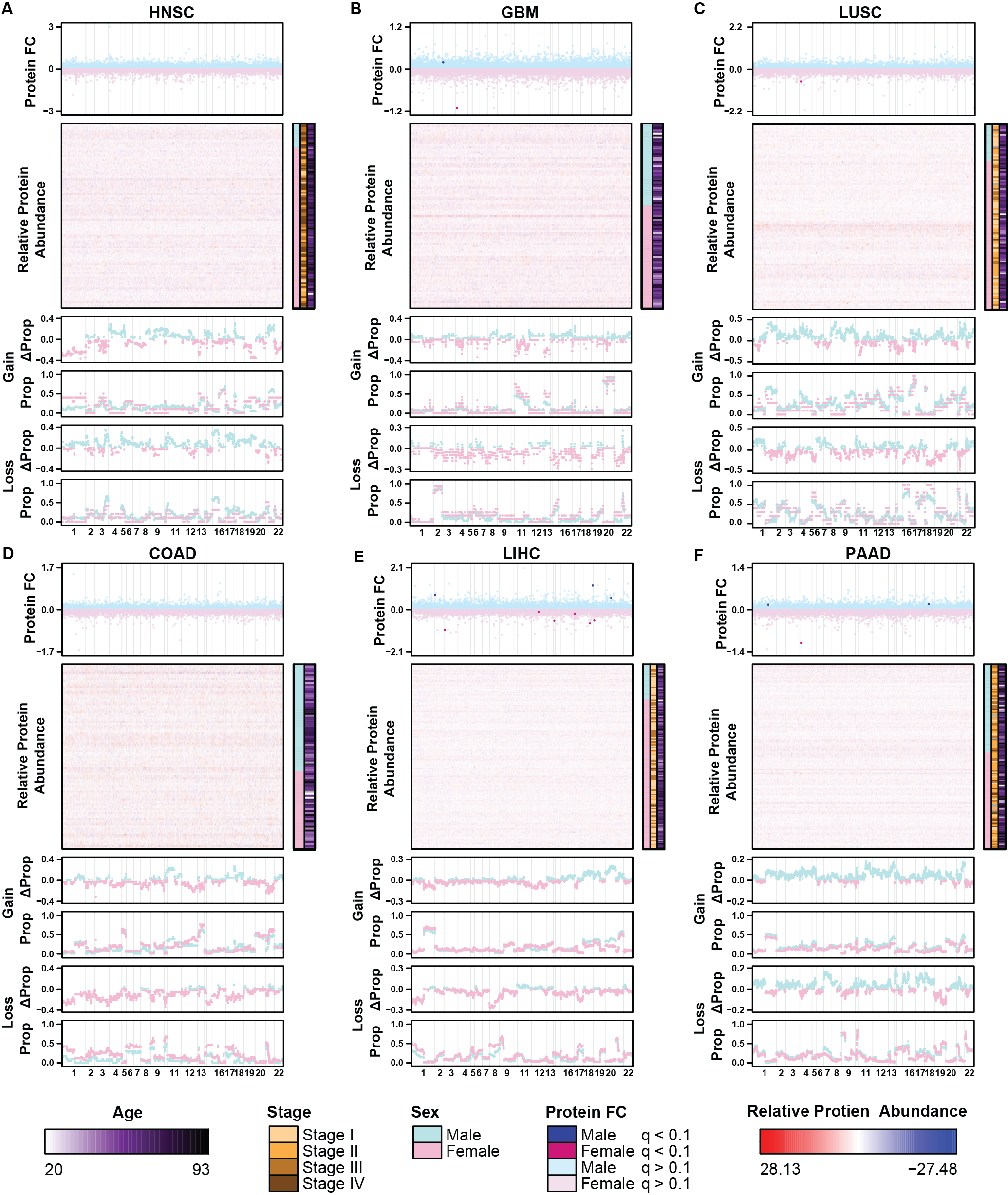

#### Supplementary Figure 5: Association of Sex Differences Between Protein Abundance and CNA

Stacked plots showing sex differences in protein abundance and copy number aberrations (CNA) across six cancer types: head and neck squamous cell carcinoma (HNSC; **A**), glioblastoma (GBM; **B**), lung squamous cell carcinoma (LUSC; **C**), column adenocarcinoma (COAD; **D**), liver hepatocellular cancer (LIHC; **E**), pancreatic adenocarcinoma (PAAD; **F**). Each panel displays, from top to bottom: fold change (FC) in protein abundance (males - females), heatmap of protein relative abundance, sex differences of copy number gain (males - females), proportion value of copy number gain in males (blue) and females (pink) separately, sex differences of copy number loss (males – females), and proportion value of copy number loss in males (blue) and females (pink). Stacked figures are aligned according to the gene coordinates in the human genome. P-values were calculated using the Mann-Whitney U test to compare the effect sizes of genes with sex-differential CNA gain or loss against those without.

### Supplementary Tables

#### Supplementary Table 1: Cancer Types, Sample Characteristics, and Summary of Sex-Differential Molecular Features

The cancer types included in the study, their abbreviations, the number of patients, tumours, normal adjacent tissues (NATs) analyzed, confounders adjusted in multivariable linear regression, the number of genes with putative sex-differential CNA status, and the number of significant sex-differential proteins identified in NATs and tumours.

#### Supplementary Table 2: Statistical Analysis Results for Proteomic Sex Differences Across Eight Cancer Types.

Statistical coefficients, standard error, p-value and q-value from univariable t-tests and multivariable linear regression for sex differences in protein abundance for each gene across eight cancer types.

#### Supplementary Table 3: Statistical Analysis Results for Sex-Differential Copy Number Aberrations Across Eight Cancer Types.

Proportion of tumours harbouring copy number aberrations (CNAs) in each sex, proportion test p-value and confidence intervals for all genes tested across eight cancer types. Gene information, including gene symbol, Ensembl ID, chromosome and position, is also included.
